## Supporting Information for "Influence of 8-oxoguanine on the thermal stability of G-quadruplex DNA and the interactions with the FANCJ AKKQ peptide"

### 8-Oxoguanine Disrupts G-Quadruplex DNA Stability and Modulates FANCJ AKKQ Binding

**CD Spectroscopy (AKKQ Secondary Structure)**

Circular dichroism (CD) data was collected on a JASCO J-815 spectropolarimeter (JASCO Inc.; Easton, MD, USA) equipped with a PTC-423S peltier system. Protein samples were analyzed either in trifluoroethanol-containing buffer (20 mM boric acid pH 7.5, 150 mM KCl or NaCl, 5% (v/v) glycerol, and 20% (v/v) trifluoroethanol) or buffer without trifluoroethanol (20 mM boric acid pH 7.5, 150 mM KCl or NaCl, and 5% (v/v) glycerol). The presence of trifluoroethanol was used to evaluate differences in protein secondary structure based on the corresponding salt condition. Spectra of FANCJ AKKQ were collected from 260 nm to 200 nm at 25 °C (S1 Fig.). Five traces were collected for each sample, and reference scans of buffer without protein were subtracted from the averaged CD signal.

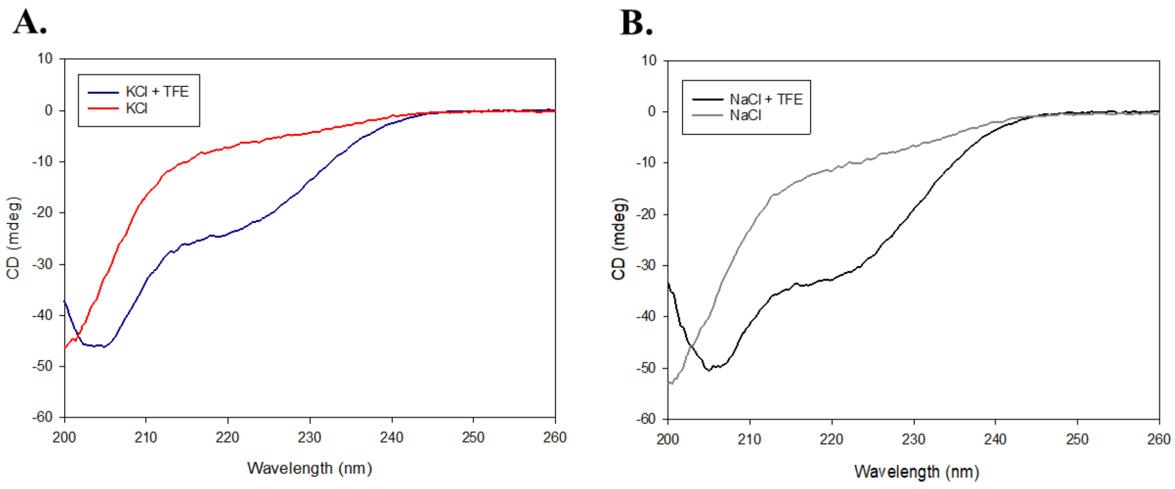


**S1 Fig. Experiments performed in the presence and absence of 20% (v/v) trifluoroethanol.** (A) CD spectra of FANCJ AKKQ taken from 260 to 200 nm in the presence (blue) and absence (red) of trifluoroethanol in 150 mM KCl. (B) CD Spectra of

FANCJ AKKQ taken from 260 to 200 nm in the presence (black) and absence (gray) of trifluoroethanol in 150 mM NaCl. Upon the addition of trifluoroethanol in either salt condition, an alpha helical conformation appears.
